# Derivation of the Relationship Between the Mass and Lifespan of Living Organisms

**DOI:** 10.1101/2022.05.09.491161

**Authors:** Anatolii Shyian

## Abstract

The relation *p=1-a* between exponents, by which the dependencies of lifespan *p* and growth intensity *a* on masses of living organisms are described, is obtained and testified.

## Introduction

The problems associated with the life span of living organisms have clearly expressed both fundamental and applied aspects. Among the first, one can single out a range of tasks related to identifying the causes that limit the life of animals. Among the second is the importance of such a parameter for applied problems is primarily due to the fact that the productivity of living organisms can be expressed in terms of their lifespan. For example, for aquatic organisms, this dependence has the form *C=kT^-n^*, where *C* is the specific productivity of animals, *T* is their duration. life, *k*>0 and *n*>0 - coefficients [1].

Recently it has been shown [2] on a significant amount of material that a close correlation between the lifespan *T* and weight *m* for a large number of species of living organisms, both plants and animals is existed. However, the reasons leading to this ratio are still unclear.

In [3] a hypothesis is put forward, “Fast versus slow aging may depend on whether the organism “grows fast” or “develops longer”: first case should be associated with high MTOR” (MTOR = Mechanistic (formally, mammalian) Target of Rapamycin). At the same time, it is noted that more massive biological organisms grow more slowly, and therefore their life expectancy is longer.

The article [4] links lifespan to the rate of telomere shortening. The question arises whether it is possible to relate the rate of telomere shortening to the exponent *a* for the phase of active growth of a biological organism. For example, this may be the case when the length of telomeres is shortened as a result of cell division. If the answer to this question is yes, then it will be possible to relate cellular characteristics to the characteristics of the entire population of organisms. This may also be important for describing a number of environmental problems.

This paper is devoted to a method for quantitatively deriving the relationship between the lifespan of living organisms and their mass.

## Theory

We will proceed from the equation accepted in modern biology for the dynamics of the individual mass of living organisms, written in the form

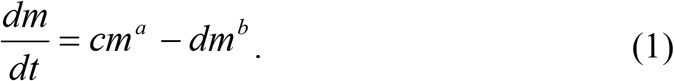

Equation (1) is usually called the Bertalanffy equation; see [1] for its derivation and interpretation.

Equation (1) allows us to introduce the concept of “definitive” mass of the considered living organisms *m_0_*, which is found from the stationary solution (1) as *m_0_=(c/d)^1/(b-a)^*. Obviously, the definitive value of the mass m0 is achievable only for *b*>*a* – we will assume that this relation is satisfied in what follows.

In general, an equation of the form

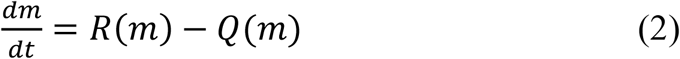

often used to describe the self-organization of open systems [5,6]. In this case, *R*(*m*) describes an increase in the exponent m due to self-organization processes, and *Q*(*m*) describes a decrease in the exponent m due to the decay processes of the biological object. In this case, there is only one point m0 at which *R*(*m_0_*)=*Q*(*m_0_*), for which *R*(*m*)>*Q*(*m*) for *m*<*m_0_* and *R*(*m*)<*Q*(*m*) for *m*>*m_0_*.

The explicit form of *R*(*m*) and *Q*(*m*) can be determined from experiment. In particular, at the initial stage of life, when the biological organism is growing rapidly, the dynamics of mass change is set mainly by *R*(*m*).

The variable m often varies within large limits, amounting to several orders of magnitude. In this case, as a rule, when processing the corresponding experimental data, doubly logarithmic coordinates are used. The data in them is processed, as a rule, using the least squares method (approximation by a linear dependence).

As a result, one obtains, for example, the following expression for *R*(*m*).

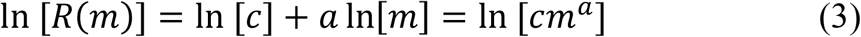

Similarly, the expression for the second term in (1) is obtained. It is in this way that scaling [7] often comes into biology.

Since, as a rule, the “initial” (e.g., after birth for mammals) and definitive masses for living organisms are related by the relation *m_ini_*<<*m_0_* [1], the characteristic time *τ_0_* required to reach the definitive mass can be estimated as

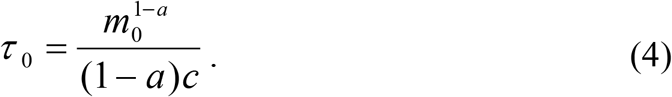

In fact, the time *τ_0_* is a characteristic (but not complete) growth time of the living organisms under consideration, because it is this time that their most intensive growth lasts.

Let the following assumption (A) be accepted:

- The lifespan *T* for living organisms is directly proportional to the characteristic time *τ_0_* required to reach their definitive mass.

In this case, the following relation will be arrived.

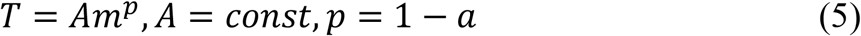

Assumption (A) actually means that the natural discrete value for measuring the lifespan *T* of living organisms is the characteristic duration of the stage of their ontogeny *τ_0_*. This is supported by the fact that the characteristic periods of the initial physiological cycles of mammals and birds can be represented in the form (5) with an indicator that practically coincides with *p* [7, 8].

It is also interesting that the assumption about the relationship between the dependence between the life span of eukaryotes and the mass of living organisms, expressed in [2], finds its direct confirmation in formula (3).

Assumption (A) also leads to this relation for the pre-exponential factors

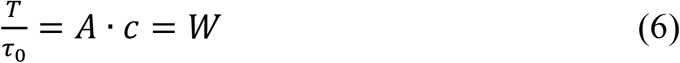

Here, in the general case, *W* may depend on the type of living organisms. However, the definition of a large group of living organisms can be carried out under the assumption *W*=const. Relation (4) can be naturally identified with the number of complete cycles of “renewal” of the body in the process of life. Such an interpretation provides ample opportunities for matching the interpretation proposed above with existing models for aging mechanisms (see [9]) and allows us to propose new quantitative approaches for processing experimental data.

It is interesting that the *W* value from relation (6) can be associated with the “characteristic” (maximum?!) number of divisions that can be performed by each cell during the life of living organisms after they reach their definitive sizes. This allows us to look from a new point of view both at the process of evolution of prokaryotes into eukaryotes, and at the processes of self-organization of multicellular organisms (and, possibly, at the genetic prerequisites for this).

## Testing

The available experimental information on the quantitative values of the parameters *a* and *p*, obtained by independent methods, makes it possible to verify the derived relation (5). As one can be seen from the Table, the results of this paper are confirmed both for a wide range of living organisms in general and for individual biological species.

**Table.**
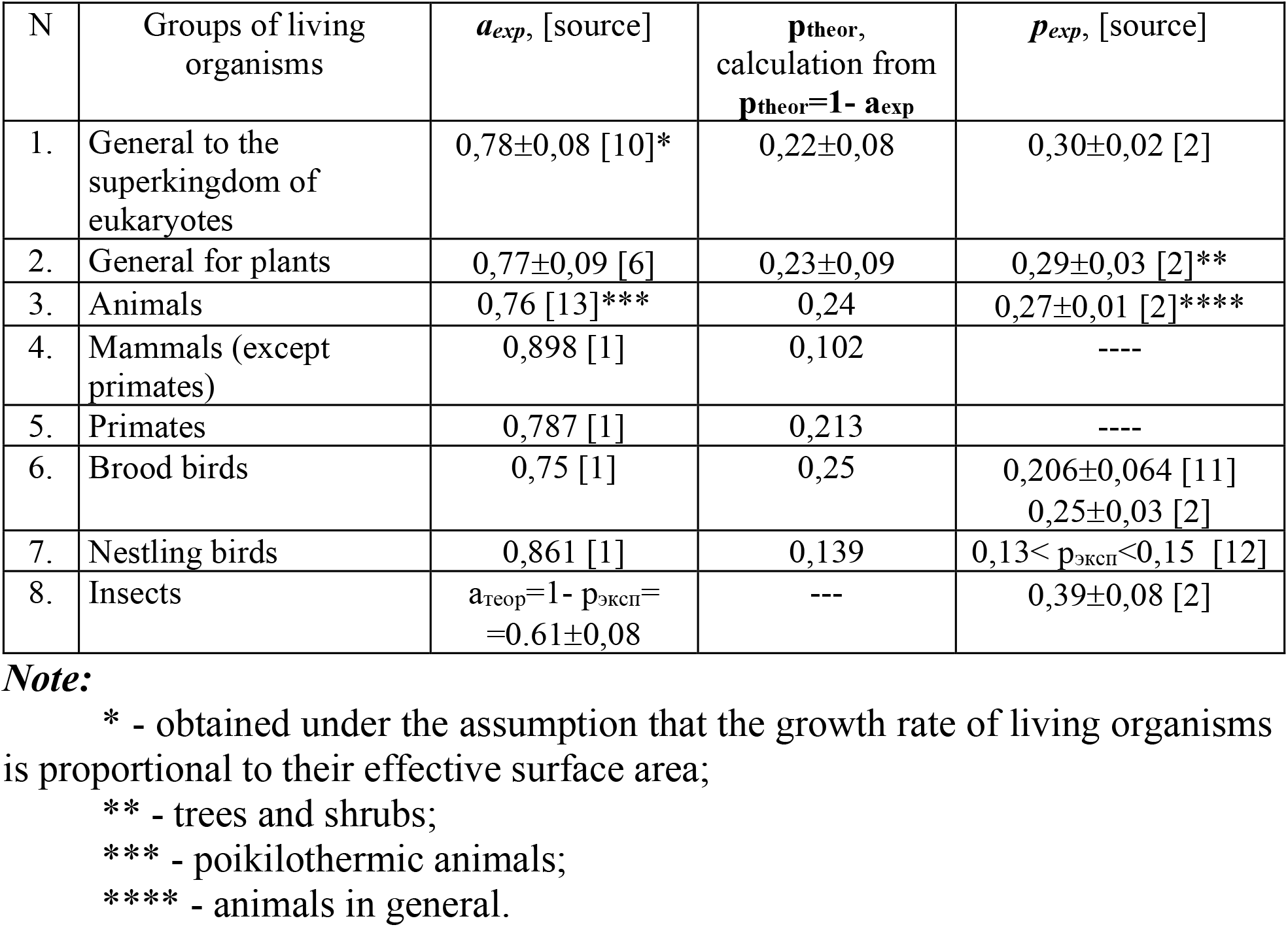
Testing the fulfillment of the relation p=1-a from (3).

The author used the databases available to him for parameters *a* and *p*. It probably makes sense to use modern databases to test relation (5).

## Conclusion

Thus, relations (5) - (6), which convey the average trend in the behavior of characteristics, can be used as the basis for the development of new methods for setting up experiments for a wide range of biological problems with different directions (from fundamental to applied), as well as methods for quantitative interpretation of the obtained results.

## Acknowledgment

Author thanks to V.E. Zaika, K.M. Khailov, N.P. Malyuta and A.V. Prazukin for helpful discussions, and O.N. Baeva for patience.

## Notes

### Competing Interest Statement

The authors have declared no competing interest.

## References

1. Zaika V.E. (1983). Comparative productivity of hydrobionts. Kyiv: Naukova Dumka. (In Russian).

2. Metelsky S.T. (1995). Relationship between the life span of eukaryotes (animals and plants) and some of their physical characteristics. Journal of General Biology. V.56, N 6. P.723–735. (In Russian).

3. Blagosklonny M.V. (2013). Big mice die young but large animals live longer. AGING. V. 5, N 4. P.227–233.

4. Whittemore K., Vera E., Martínez-Nevado E., Sanpera C., Blasco M.A. (2019). Telomere shortening rate predicts species life span. PNAS. V. 116, N 30. P.15122–15127.

5. Shiyan A.A. (1997). The Mass distribution of Biological Systems as a Characteristic of Their Interaction with the Surrounding Medium. Biophysics. V.42. N 6. P.1173–1178.

6. Shiyan A.A. (1998). On the Problem of Elaboration of New Criteria for Control of Hierarchical Socio-Economic Systems. Journal of Automation and Information Sciences. No. 4-5. P.216–225.

7. Schmidt-Nielsen K. (1984). Scaling: why is animal size so important? Cambridge University Press.

8. Lindstedt S.L., Calder W.A. Body size, physiological time, and longevity of homeothermic animals. Quart. Rev. Biol. 1981. V.59. P.1–16.

9. Frolkis V.V., Muradyan H.K. (1988). Experimental ways to increase lifespan. Leningrad, Nauka. (In Russian).

10. Khailov K.M., Prazukin A.V., Kovardakov S.A., Rygalov V.E. (1992). Functional morphology of marine multicellular algae. Kyiv: Naukova Dumka. (In Russian).

11. Gavrilov V.V. (1995). Association of maximum lifespan with social organization in free-living waders (CHARADPII, AVES). Journal of General Biology. V.56, N5. P.529–538. (In Russian).

12. Paevsky V.A. (1985). Bird demographics. Leningrad, Nauka. (In Russian).

13. Alimov A.F. (1992). Characteristics of populations, communities of hydrobionts and mass of animals. Reports of the Russian Academy of Sciences. V.323, N3. P.588–591. (In Russian).

